# Learning to See Like a Child: Why Viewpoint Diversity is Fundamental for Human-Aligned Object Recognition

**DOI:** 10.64898/2026.05.27.727784

**Authors:** Yifan Luo, Niklas Müller, H. Steven Scholte

**Affiliations:** University of Amsterdam; KU Leuven

**Keywords:** convolutional neural networks, object representation, viewpoint diversity, out-of-distribution generalization, synthetic dataset

## Abstract

Deep convolutional neural networks match human accuracy on standard object recognition tasks but fail to recognize familiar objects from novel viewpoints. Humans, however, develop viewpoint-invariant recognition at an early age through diverse visual experience. This gap in visual experience may explain why models diverge from humans in object recognition. Holding dataset size constant, we show that greater viewpoint diversity substantially improves generalization to novel views. Using a synthetic 3D dataset with systematically controlled viewpoints, we reveal a core trade-off: restricted-view training yields rapid learning and near-ceiling in-distribution accuracy but collapses on held-out viewpoints, whereas viewpoint-diverse training learns more gradually yet generalizes robustly. Increasing viewpoint diversity disrupts texture regularities while preserving global shape, driving networks to prioritize shape over texture - the same strategy that underlies human object recognition. Partitioned Grad-CAM analyses further show that viewpoint-diverse models maintain object-centered attention. These findings parallel developmental accounts of multi-view learning and identify viewpoint diversity as an important factor for robust, human-aligned vision.

## Introduction

Convolutional neural networks (CNNs) have achieved remarkable success in object recognition and are widely studied as computational models of human vision He et al. (2016); Kriegeskorte (2015). Yet, unlike humans, they struggle to recognize objects from novel viewpoints, particularly from the non-canonical perspectives - unusual angles that are rarely represented in training datasets and often require real-world exploration or 3D simulation to capture Madan et al. (2022); Sakai et al. (2022); Alcorn et al. (2019); Barbu et al. (2019). For example, recognizing a mug from the frontal side view is straightforward, but identifying it from below can pose a significant challenge for CNNs. Humans, too, are slower and less accurate with such perspectives Tarr (1995); Milivojevic (2012), yet remain markedly better than CNNs at generalizing across viewpoints Ollikka et al. (2025). This limitation hinders CNN deployment in real-world applications and raises concerns about their plausibility as models of human visual processing.

One reason for this gap lies in differences between human visual experience and the training dataset of CNNs. CNN performance is tightly linked to the composition of training datasets: widely-used image datasets such as Caltech 101 Fei-Fei et al. (2004) and ImageNet Deng et al. (2009) provide broad category coverage but little systematic variation in viewpoint Barbu et al. (2019). In contrast, humans or even non-human animals can develop viewpoint-invariant recognition through repeated exposure to objects from many perspectives early in life Tarr (1995); Milivojevic (2012); Wood and Wood (2016). This contrast highlights a fundamental mismatch in viewpoint diversity between traditional CNN training regimes and human vision development. Here, we try to bridge this gap by training on viewpoint-diverse stimulus sets.

Another notable characteristic of CNNs is their reliance on visual cues that differ from those typically used in human vision. A well-known example is texture bias – the tendency to prioritize local textural features over shape information Geirhos et al. (2019). This contrasts sharply with the shape-dominant processing observed in human perception Landau et al. (1988); Orhan and Lake (2024); Huber et al. (2023); Cutzu and Edelman (1998); Diesendruck and Bloom (2003), a strategy thought to emerge from extensive multi-viewpoint experience with objects during development Tarr (1995); Kraebel and Gerhardstein (2006); Wood and Wood (2016); Müller et al. (2025). CNNs are also prone to exploit background context for classification, in particular shallow networks benefit from congruent object–background associations and fail in incongruent ones, indicating that background information leaks into object representations Seijdel et al. (2020). More broadly, CNNs can sometimes classify objects from background alone Zhu et al. (2017), or misclassify when incongruent scene elements are introduced Rosenfeld et al. (2018). These findings underscore that CNNs, unlike humans, do not reliably separate objects from their content and struggle with figure-ground segmentation.

We draw inspiration from natural viewpoint diversity in human visual development and leverage 3D object image synthesis to systematically examine how viewpoint diversity in training data influences CNN performance and alignment with human behavior. Since viewpoint variation disrupts texture consistency while preserving shape cues, training with greater viewpoint diversity may help counteract CNNs’ over-reliance on texture or dependence on background context. While our approach does not capture the full dynamics of human perception, it allows us to isolate whether viewpoint diversity as a fundamental aspect of developmental visual experience improves CNN object recognition.

We first investigate how controlled viewpoint diversity affects generalization using synthetically rendered datasets of common objects that vary in their degree of viewpoint diversity. Specifically, we ask whether restricted-view training produces brittle, view-specific object recognition solutions that perform well on familiar images but collapse when objects are seen from new perspectives, whereas diverse training fosters representations that generalize more robustly.

To disentangle viewpoint generalization from robustness to low-level corruption, we further evaluate models on corrupted versions of the synthesized images Hendrycks and Dietterich (2019). This allows us to test whether viewpoint diversity supports invariance beyond pixel-level alterations – specifically, whether it also stabilizes recognition under viewpoint change, in ways that standard data augmentation might not capture. In this way, we ask whether viewpoint diversity induces qualitatively different invariances which are anchored in object structure rather than low-level noise robustness.

Finally, we examine the spatial focus of model activations using Gradient-weighted Class Activation Mapping (Grad-CAM; selvaraju2017gradcam, chattopad-hay2018gradcam++). By leveraging ground-truth background segmentation in our synthetic datasets, we decompose activation maps into object, border, and background components, enabling us to ask how viewpoint diversity affects the stability and object-centeredness of model attention. This analysis directly probes the segmentation problem: restricted-view models are known to use information from background regions, whereas humans exhibit object-based attention Egly et al. (1994); Chen (2012); Lindsay (2020). The key question is whether viewpoint-diverse models, trained across varied perspectives, maintain more stable, object-centered focus under novel views.

Together, these questions motivate an investigation into how viewpoint diversity shapes the robustness, generalizability, behavioral human-alignment, and object representations in CNNs. Further, by probing models with controlled synthetic datasets, we will evaluate to what degree systematic exposure to viewpoint diversity can mitigate texture reliance and promote object-centered representations.

## Methods

### Data Synthesis

We constructed a synthetic dataset based on the 3D objects from Objaverse 1.0 dataset Deitke et al. (2023). We selected 1,544 unique object instances spanning 32 object categories (see Figure S1). All 3D objects were integrated into a Unity environment (version 2022.3.15f1c1; unity) to enable controlled rendering. Objects were randomly instantiated within a fixed spatial range, and a virtual camera was positioned to capture images from systematically varied viewpoints. To prevent networks from learning spurious associations between specific backgrounds and object categories, each object was rendered across four distinct virtual environments (meadow, forest, desert, and industrial), ensuring that object identity was independent of scene context Seijdel et al. (2020).

To investigate the effect of viewpoint diversity on CNN training, we generated four training datasets with increasing levels of viewpoint variation (see Figure 1). The viewpoint ranges were defined by the polar (*θ*) and azimuth angle (*ϕ*) of the camera origin. The viewpoint diversity levels were defined as **fixed** (*θ* = *π, ϕ* = 0.5*π*), **extra restricted** (*θ* ~*U* (1.4*π*, 2*π*), *ϕ* ~ *U* (0.4*π*, 0.8*π*)), **restricted** (*θ* ~ *U* (0.8*π*, 2*π*), *ϕ* ~ *U* (0.2*π*, 0.8*π*)), and **full** (*θ* ~ *U* (0, 2*π*), *ϕ* ~ *U* (0, *π*)). These were constructed by increasing the range of camera placement around the object. In all cases, the camera was positioned at a fixed radius of 1 unit from the object center, with a random positional offset of 0 to 0.1 units applied to the camera’s aim vector to simulate minor perspective jitter. Each object was rendered in 30 unique viewpoints per scene at a resolution of 256 × 256 pixels. To normalize apparent object scale across images, all 3D objects were automatically re-scaled such that the longest side of their bounding box fell within a standardized range of 0.7 to 1.0 Unity unit lengths.

**Figure 1.**
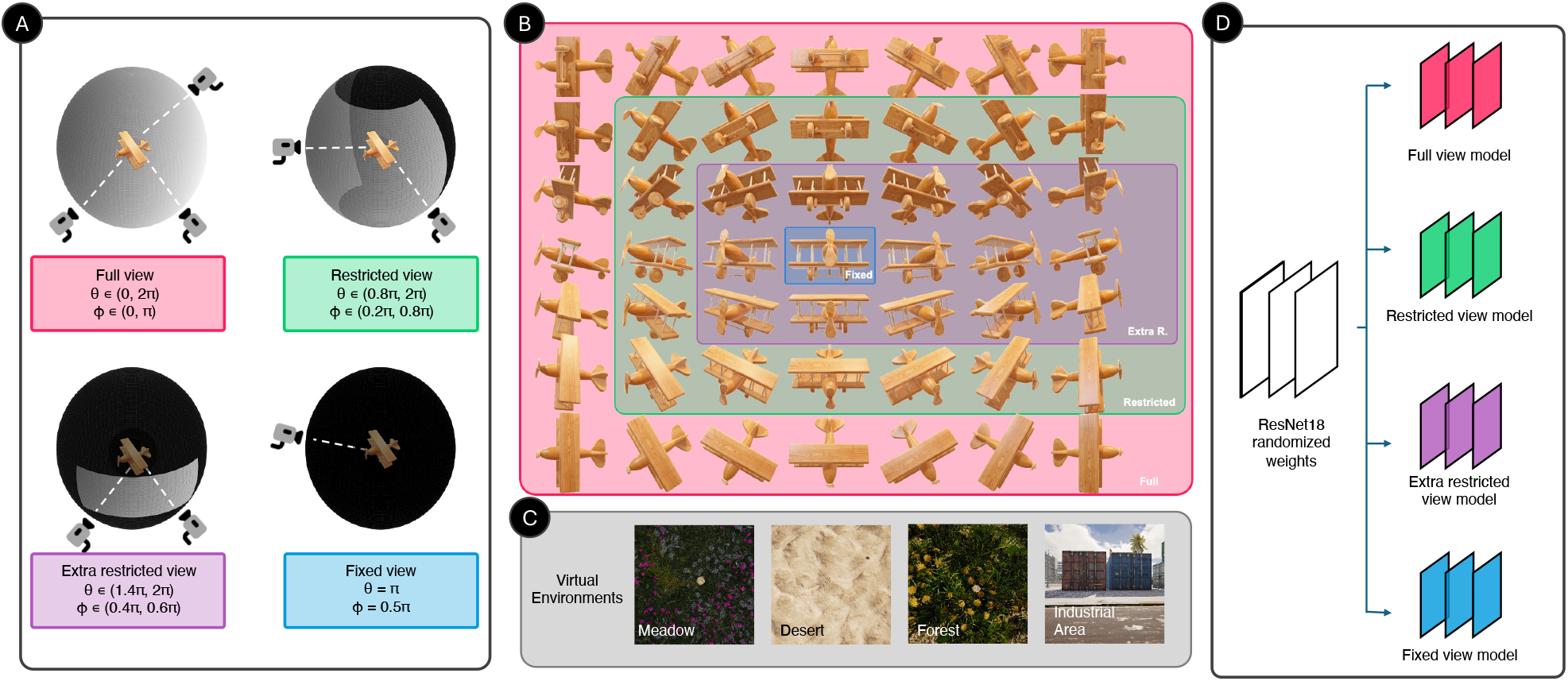
Overview of dataset synthesis with controlled viewpoint diversity and model training. **A. Schematic demonstration of viewpoint manipulation.** Objects were rendered from systematically varied viewpoints by positioning a virtual camera on a viewing sphere around the object (camera icons represent possible placements). Four viewing conditions (full, restricted, extra restricted, fixed) were defined by polar angle (*θ*) and azimuth angle (*ϕ*) ranges. These ranges determined the diversity of viewpoints included in each dataset. **B. Comparison of viewpoint diversity**. Example images of a wooden toy airplane illustrate the nested relationship among the four viewpoint conditions. The full-view dataset (red box) encompasses the widest range of perspectives, within which restricted (green box), extra restricted (purple box), and fixed (blue box) conditions represent progressively narrower subsets of orientations. This design ensures that all datasets contain the same object categories but differ systematically in viewpoint diversity. **C. 3D environments used as backgrounds**. Each object was rendered across four distinct virtual environments: meadow, desert, forest, and industrial area. **D. Training Diet**. Four separate ResNet-18 models were trained from randomized weights using datasets with varied viewpoint diversity.

Each training dataset contained approximately 185,000 images. In-distribution viewpoint (IDV) test sets were generated using the same sampling strategy as the corresponding training sets, resulting in four additional test datasets of approximately 61,000 images each. IDV test sets also serve as bases for subsequent alterations involving style transfer and image corruption. To evaluate held-out viewpoint (HOV) generalization, we also constructed a complementary viewpoint dataset (61,000 images) that samples from camera angles not captured by the restricted-view set, extra restricted-view set, or fixed-view set. However, it should be explicitly noted that by design, no viewpoint is HOV with respect to the full-view dataset. For completeness and comparative analysis, we also tested models trained on full-view set against the complementary viewpoint set. It is worth noting that in this study, we define IDV and HOV exclusively in terms of viewpoint distribution, allowing us to isolate and quantify the effect of viewpoint diversity on learned object representations.

### Training Scheme

We trained separate instances of the ResNet-18 architecture He et al. (2016) from scratch and on the four datasets corresponding to different levels of view-point diversity. For each viewpoint level, we repeatedly trained 5 model instances using different random seed. We used ResNet-18 because it provides a good balance between representational capacity and computational efficiency, enabling systematic exploration of viewpoint diversity effects across multiple conditions while maintaining comparability with established benchmarks.

The final fully connected layer was modified to output logits over 32 object categories. All models were trained on an NVIDIA Tesla V100-PCIE-32GB GPU (CUDA 12.5, driver version 555.42.06) using PyTorch. Training was performed from scratch with ResNet-18 backbones for 30 epochs, a batch size of 32, and stochastic gradient descent (learning rate = 0.001, momentum = 0.9, weight decay = 0.001) with StepLR scheduling (step size = 10, *γ* = 0.1). The loss function was cross-entropy. Models’ shape bias, robustness to image corruption, or Grad-CAM++ were evaluated using the final (30th) epoch checkpoint unless stated otherwise. Input images were first center-cropped to 224 × 224 pixels to ensure consistent input dimensions, after which the object remains visible and intact. Data augmentation included random horizontal flipping, and all images were normalized using the standard ImageNet mean and standard deviation. Each dataset was partitioned into 80% training and 20% validation subsets using stratified sampling to maintain class balance. For each epoch, top-1 training and validation accuracy were recorded. After training, the saved models were evaluated on both their corresponding IDV test set and the complementary HOV viewpoint dataset to assess generalization (see Figure 2).

**Figure 2.**
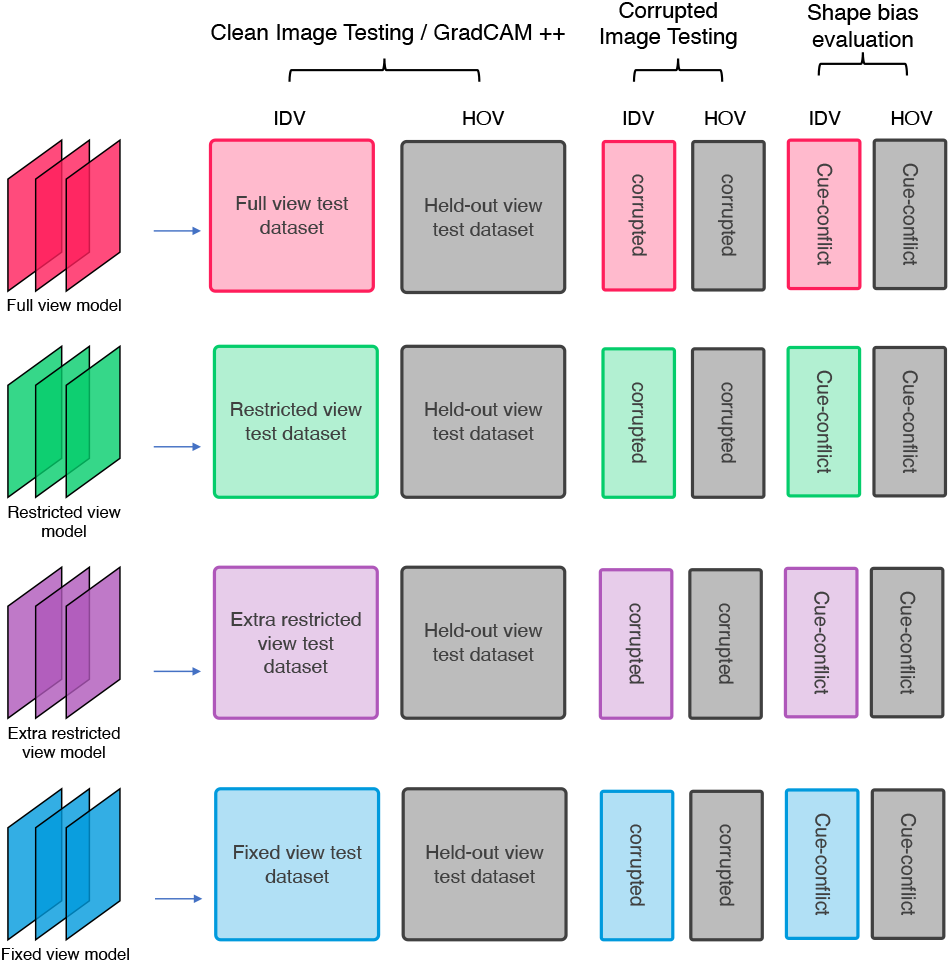
Overview of model evaluation pipeline. Trained models were evaluated across three testing streams: (1) *Clean image testing and Grad-CAM++ analysis*, using independently generated IDV (colored boxes) and HOV (gray boxes) test image sets; (2) *Corrupted image testing*, using a balanced subsample of IDV and HOV test sets which were processed with 19 corruption types at five severity levels; and (3) *Shape bias evaluation*, using a balanced subsample of IDV and HOV test sets which were processed with style-transfer.

**Figure 3.**
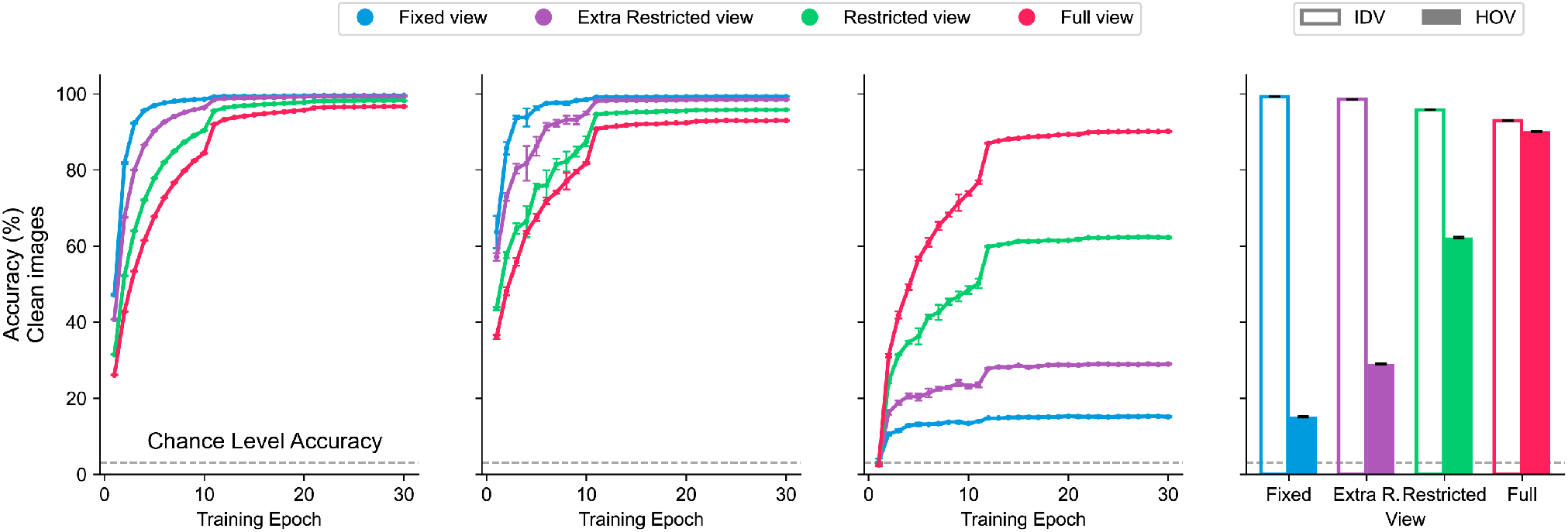
Viewpoint diversity improves HOV generalization at the cost of IDV accuracy. Training accuracy (far left), IDV (in-distribution viewpoints) test accuracy (middle left), and HOV (held-out viewpoints) test accuracy (middle right) are plotted across 30 training epochs for models trained with different levels of viewpoint diversity (fixed, extra restricted, restricted, full). The far right bar plot summarizes final-epoch IDV and HOV accuracy to show the trade-off between near-perfect IDV performance in viewpoint-impoverished models and higher HOV generalization in viewpoint-diverse models. Error bars indicate the standard error of accuracy for each data point. The dashed line in each plot indicates chance accuracy in object recognition task.

### Image Corruption

We evaluated model robustness under low-level visual perturbations using the image corruption benchmark by hendrycks2018benchmarking. From each of the five test datasets, we randomly sampled 640 images while maintaining a balanced distribution across object categories and background types to match similar benchmarks (Tiny ImageNet-C, hendrycks2018benchmarking). Each image was then processed with 19 types of common corruptions (e.g., Gaussian noise, motion blur, brightness, frost, etc.) across five severity levels (see Figure 4). The resulting corrupted datasets were used to evaluate each model on its corresponding IDV and HOV test sets.

**Figure 4.**
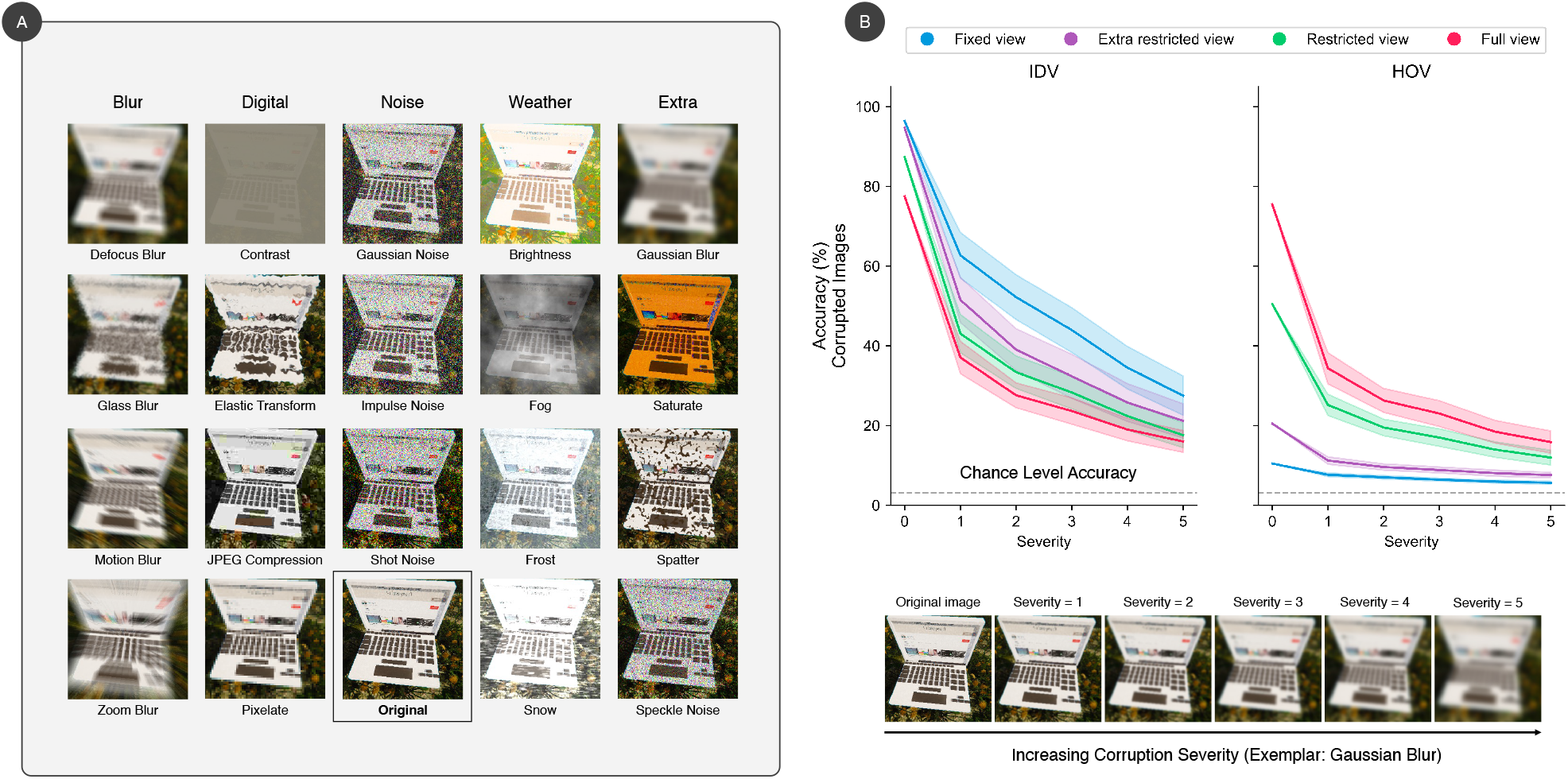
Viewpoint diversity enhances corruption robustness on HOV viewpoints, while slightly reducing robustness on IDV data. **A. Image corruption methods.** Test images were processed with 19 common corruption types spanning blur, digital, noise, weather, and additional distortions Hendrycks and Dietterich (2019). Each corruption was applied at five severity levels, illustrated here with a laptop example. **B. Categorization accuracy on corrupted test images**. Accuracy is plotted as a function of corruption severity (levels 0–5), averaged across the 19 corruption types. The left panel shows accuracy on IDV viewpoints, while the right panel shows accuracy on HOV viewpoints. Lines denote mean accuracy across models, and shaded areas represent standard error. Dashed horizontal lines indicate chance-level accuracy.

### Style Transfer and Texture–Shape Bias

To examine the texture–shape bias in model predictions, we employed neural style transfer Gatys et al. (2016) to create cue-conflict stimuli. In these images, the object shape is preserved from a content image, while the texture is replaced with that from a different object class (see Figure 5). For each of the five test datasets, we randomly sampled 512 content images balanced across categories and backgrounds. Each content image was then stylized (5,000 optimization steps, based on VGG-19 feature extractor) using textures from four randomly selected different object categories, resulting in five cue-conflict datasets, each comprising 2,560 images.

**Figure 5.**
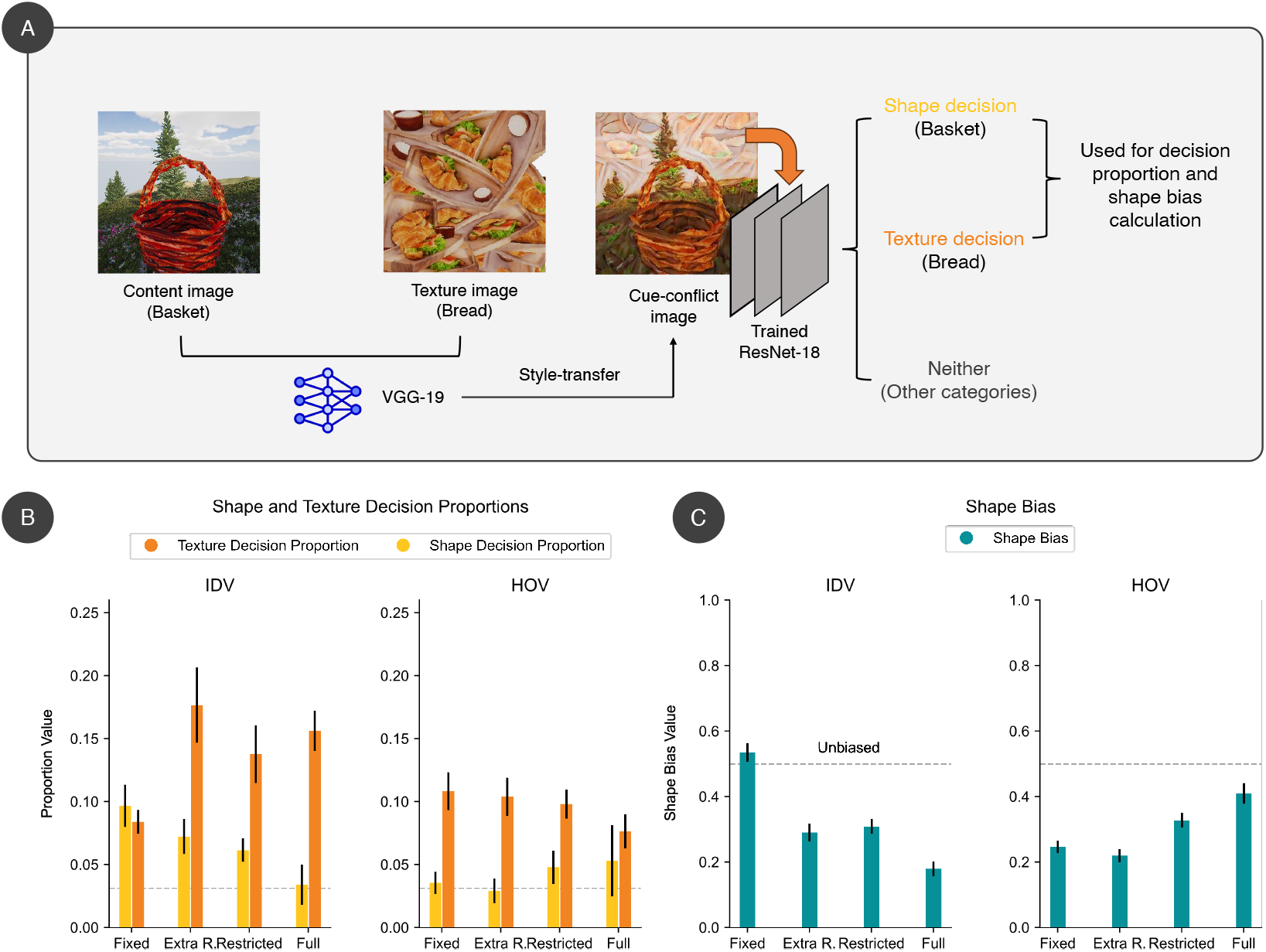
Viewpoint diversity increases shape bias on HOV data but reduces it on IDV data. **A. Cue-conflict stimuli generation.** Stimuli were generated by combining the shape from a content image with the texture of an unrelated image using neural style transfer. Model predictions were categorized as shape-based (aligned with the content image), texture-based (aligned with the style image), or neither. **B. Shape and texture decision proportions**. The proportions of shape-based (yellow) and texture-based (orange) decisions on cue-conflict images, separately for IDV (left) and HOV (right) test sets. Dashed horizontal lines mark proportion at chance level. **C. Shape bias**. Models trained on viewpoint-diverse datasets show reduced shape bias on familiar IDV views (left) but increased shape bias on novel HOV views (right). Dashed horizontal lines mark unbiased preference. Error bars represent standard error.

Models were evaluated on both the IDV and HOV cue-conflict datasets. For each image, a prediction matching the original content category was defined as a *shape decision*, whereas a prediction matching the style source was considered a *texture decision*. The *shape bias* was quantified as the proportion of shape decisions relative to the total number of shape and texture decisions:

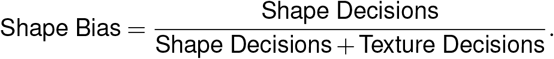

### Partitioned Grad-CAM

To gain insights into model decision processes, we employed Grad-CAM++ Selvaraju et al. (2017); Chattopadhay et al. (2018) to visualize spatial attention on both IDV and HOV images, including those with corruption and style transfer (Figure 6). Grad-CAM++ maps were computed using the final convolutional block of ResNet-18 as the target layer. Grad-CAM++ highlighted image regions that contribute most strongly to the model’s predicted label.

**Figure 6.**
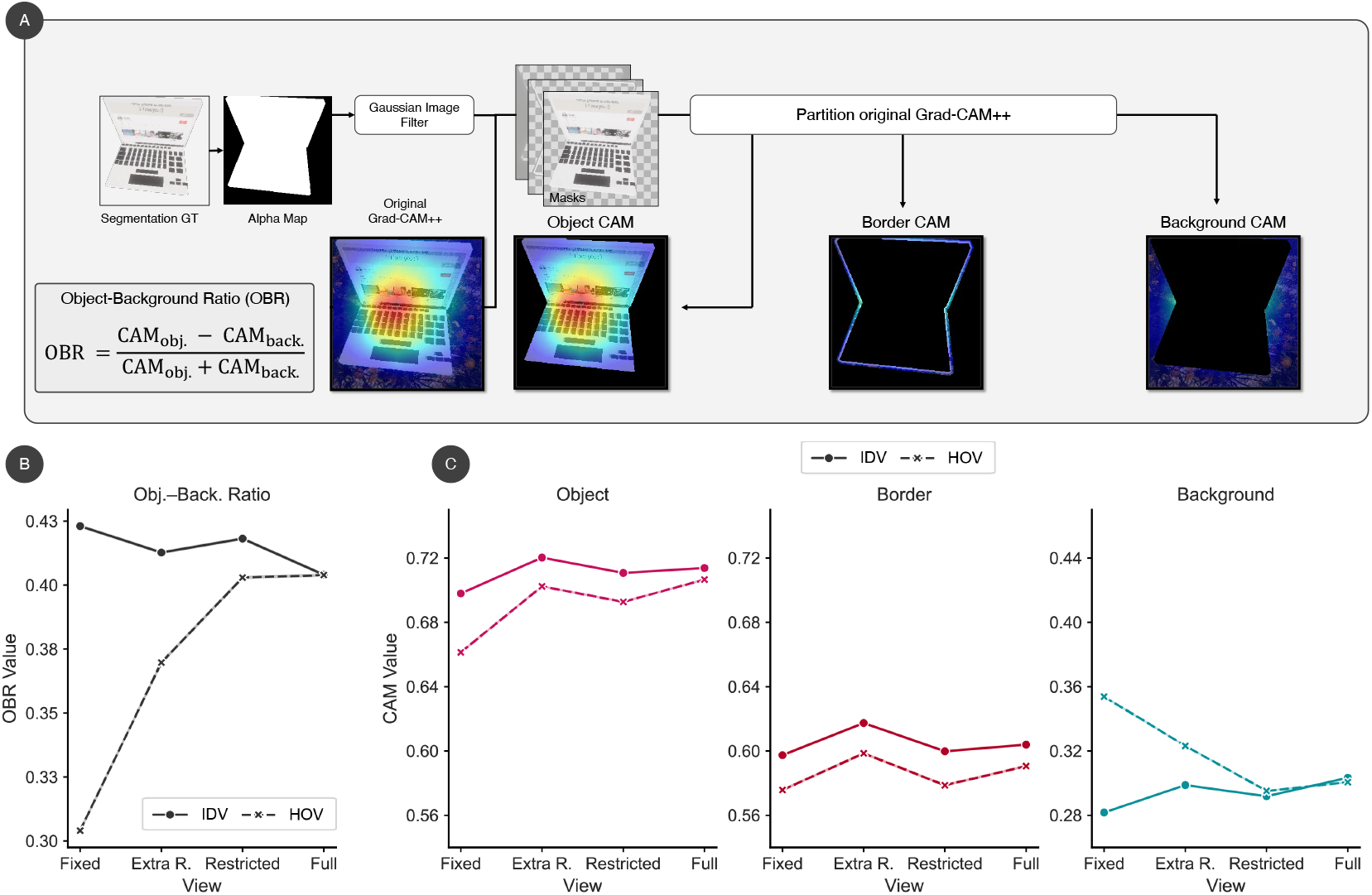
Viewpoint diversity improves spatial stability of model attention under distribution shift. **A. GradCAM maps.** To analyze model attention, Grad-CAM++ Chattopadhay et al. (2018) heatmaps were computed for each test image and partitioned using ground-truth object segmentation. A Gaussian-filtered alpha mask was used to separate activations into object, border, and background regions. Values are averaged for IDV (solid lines) and HOV (dashed lines) images across models trained with different viewpoint diversities. **Object–Background Ratio (OBR)**. The OBR captures the extent to which attention remains object-focused rather than background-biased. Higher OBR indicates more stable object-centered attention under viewpoint shift. **C. Grad-CAM values in object, border, and background divisions**.

To further dissect the spatial distribution of model attention, we partitioned each Grad-CAM heatmap using pixel-level object segmentation ground truth. This was obtained by re-rendering all images in Unity under identical object and camera configurations but with a transparent background. The alpha channel from these images yielded binary segmentation masks, with fore-ground object pixels set to 1 and background pixels set to 0. A Gaussian filter (*σ*=1 pixel) was applied to the alpha mask to create soft transitions at object borders. We then separated the Grad-CAM into three components:

1. **Object CAM:** Regions where the filtered alpha value equals 1, representing attention on the object itself.
2. **Background CAM:** Regions where the filtered alpha value equals 0, representing attention outside the object.
3. **Border CAM:** Regions where the filtered alpha value lies between 0 and 1, capturing attention at object boundaries.

This partitioning allowed us to quantify how viewpoint diversity influences the spatial localization of model attention.

We evaluated the partitioned CAM values for each model on both IDV and HOV test datasets. For each image, we computed a Grad-CAM++ activation map corresponding to the model’s predicted label. This resulted in a single-channel heatmap where each pixel held an activation value between 0 and 1, indicating its contribution to the final decision.

Using the filtered alpha map, we then averaged the Grad-CAM values separately within each of the three partitions (object, border, and background). This yielded per-image scalar metrics for object CAM, border CAM, and background CAM. These values were then averaged across all images in each test condition to quantify how model attention was distributed spatially and how this distribution varied as a function of viewpoint diversity. By comparing CAM partition scores across models and test types, we assessed the degree to which viewpoint-diverse training encourages object-focused attention and reduces reliance on background or border regions under viewpoint distributional shifts. To directly quantify the relative concentration of model attention on the object versus the background, we define a normalized metric Object-Background Ratio (OBR),

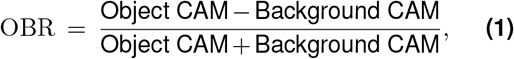

where “Object CAM” and “Background CAM” are the per-image averages of the Grad-CAM++ values within the object and background partitions, respectively. OBR lies in (− 1, 1): values > 0 indicate proportionally stronger attention on the object than the background, values < 0 indicate the opposite, and values near 0 indicate comparable allocation. Because OBR is a normalized difference, it is less sensitive to global scaling of the heatmap and complements raw partition scores by emphasizing where attention is concentrated. Positive OBR values indicate proportionally stronger object-than background-focused saliency, while lower values reflect a relative shift toward background reliance.

## Results

### Viewpoint Diversity Promotes Generalization Across Novel Viewpoints

The model performance on IDV and HOV data is summarized in Figure 3, confirming a fundamental trade-off between IDV performance and HOV generalization as a function of viewpoint diversity. To aid interpretation, we grouped the fixed and extra restricted view conditions as *viewpoint-impoverished*, and the restricted and full view conditions as *viewpoint-diverse*. Note that by design, the held-out-viewpoint (HOV) set contains camera orientations that are unseen for the fixed, extra restricted, and restricted conditions but not truly novel for the full-view condition (see Section). Models trained on viewpoint-impoverished datasets reached near-ceiling training accuracies (fixed: 99.58%, extra restricted: 99.38%) and high IDV test accuracies (fixed: 99.29%, extra restricted: 98.58%). Models trained with viewpoint-diverse data achieved slightly lower training (restricted: 98.23%, full: 96.72%) and IDV test accuracy (restricted: 95.72%, full: 92.99%).

However, this pattern reverses when evaluating HOV generalization. viewpoint-impoverished models exhibited severe performance drops on novel viewpoints, with HOV test accuracy falling below 30% (fixed: 15.62%, extra restricted: 28.93%). In contrast, viewpoint-diverse models generalized substantially better to novel viewpoints, with the full-view model achieving 90.33% and the restricted-view model reaching 62.17%.

### Viewpoint Generalization is Distinct from Corruption Robustness

Corruption tests revealed that viewpoint generalization is not simply a byproduct of robustness to image corruption. Figure 4 shows model accuracy on corrupted IDV and HOV viewpoint test sets, plotted against corruption severity. As expected, accuracy declines as corruption severity increases in all models. Crucially, the relationship between viewpoint diversity and corruption robustness reveals a clear dissociation. Models trained with fixed or extra restricted viewpoints perform well under mild corruption on IDV data (e.g., fixed-view: 62.72% at severity level 1), but collapse under HOV viewpoint corruption (7.62%). This indicates a strong reliance on view-specific cues that do not generalize when the view-point changes.

By contrast, models trained with full viewpoint diversity show more balanced performance across both IDV and HOV corrupted inputs. While their accuracy on IDV-corrupted images is somewhat lower under mild perturbations (full-view: 37.07% at severity 1), they achieve substantially better performance on corrupted HOV views (34.35%), exceeding all viewpoint-impoverished models. This dissociation demonstrates that viewpoint diversity induces structural invariance not captured by conventional augmentation.

### Viewpoint Diversity Modulates Shape Bias Based on Generalization Context

To assess the nature of the representations developed by the models in object recognition, we evaluated shape bias using cue-conflict images that disentangle shape and texture cues (Figure 5). Models trained with greater viewpoint diversity exhibited a consistent trend: shape bias decreased on IDV data but increased on HOV data. Specifically, shape bias on IDV test images declined with viewpoint diversity (fixed: 0.54, extra restricted: 0.29, restricted: 0.31, full: 0.18), while shape bias on HOV test images increased (fixed: 0.25, extra restricted: 0.22, restricted: 0.33, full: 0.41).

Analysis of decision patterns revealed why: on IDV data, restricted-view models favored texture-based choices, while on HOV data, full-view models made substantially more shape-consistent classifications. Thus, viewpoint diversity produces a shape bias that emerges specifically under viewpoint distribution shift, when generalization is required.

### Viewpoint Diversity Stabilizes Object-Centric Attention under Viewpoint Change

We next examined spatial processing with Grad-CAM++ Chattopadhay et al. (2018), partitioning activations into object, border and background regions (see Figure 6). This analysis revealed that viewpoint diversity stabilizes object-centered attention under viewpoint distribution shift.

Although all models showed reduced object activation under HOV inputs, the decline was modest in viewpoint-diverse models. Restricted-view networks exhibited a double dissociation: object activation decreased (0.70→0.66) while background activation increased (0.28→0.35). By contrast, full-view training preserved both object activation (0.71→0.71) and background stability (0.30→0.30). Importantly, these activation patterns were highly consistent across individual images (variability ≤ 0.001), indicating that the observed differences reflect stable representational strategies rather than noisy fluctuations. These results indicate that viewpoint diversity anchors attention to the object itself, preventing the leakage of activations into background regions that characterizes restricted-view training.

We further quantified the balance of attention between object and background using the OBR (Eq. 1). Across all conditions, OBR values were substantially greater than zero, confirming a consistent object bias. However, the degree of contrast varied systematically by training regime and test distribution. Restricted-view models achieved the highest OBR overall (IDV: 0.42 ± 0.0004; HOV: 0.40 ± 0.0004), reflecting strong and relatively stable object-centered saliency across distributions. By comparison, extra restricted and fixed-view models showed sharper declines under HOV evaluation (Fixed: 0.42 → 0.30; Extra-R: 0.41 → 0.37). Full-view models exhibited intermediate but balanced values (IDV: 0.40 ± 0.0004; HOV: 0.40 ± 0.0004), indicating robust object focus that generalized smoothly across viewpoint shifts without large drops.

## Discussion

Our study demonstrates that training CNNs with diverse viewpoints of objects improves both generalization and object representation quality. While viewpoint-restricted models achieve near-perfect accuracy on familiar, IDV views, they fail on novel, HOV viewpoints. In contrast, viewpoint-diverse models maintain strong performance across viewpoints, despite slightly lower IDV accuracy. Corruption experiments, shape bias evaluation, and Grad-CAM++ analysis together reveal that viewpoint-diverse models develop more object-centered attention, which is more aligned with human strategy. We observed that viewpoint-diverse models flexibly shift from texture to shape-based processing under novel views, maintain stable object-focused attention, and show reduced background interference. The converging evidence demonstrates that viewpoint diversity during training encourages models to shift from over-relying on view-specific features toward constructing representations that capture the underlying structural invariants of objects.

### Mechanistic Insights

During the model training, we showed that viewpoint-restricted models converge rapidly, reaching near-ceiling accuracy on more familiar, IDV views within a few epochs. However, this fast learning reflects reliance on view-specific patterns rather than abstraction of object structure. In contrast, viewpoint-diverse models learn more gradually, with slower initial gains but sustained improvement over training. The two kinds of trajectories suggest that diverse input increases variability in the training signal, forcing networks to progressively uncover higher-level, more transferable features. This slower and more gradual learning trajectory mirrors coarse-to-fine developmental patterns observed in infants and is consistent with computational models that simulate such developmental dynamics Smith and Slone (2017).

We demonstrated that viewpoint-restricted and viewpoint-diverse models develop qualitatively different object representations, both in terms of robustness and the cues they prioritize. Restricted models maintain accurate recognition despite corruptions on familiar views but collapsed when corruption coincided with novel viewpoints, showing their dependence on narrowly defined visual regimes. By contrast, although viewpoint-diverse models show reduced performance under corruption with familiar viewpoints, they sustained their performance under corruption compounded with novel viewpoints, suggesting that diversity drives networks to latch onto structural features that remain stable across viewpoint shifts without conferring extra robustness to low-level perturbations. These results highlight that the stability of representations depends less on resistance to pixel-level noise and more on encoding viewpoint-invariant object structure.

Shape-bias analyses further revealed that viewpoint diversity alters how networks weigh texture versus shape cues, providing a window into the representational strategies underlying this stability. Restricted-view models appeared more shape-biased on familiar IDV inputs, but this effect likely reflects overfitting: when training views are limited, texture and shape cues are tightly correlated, so reliance on superficial textures can masquerade as genuine shape sensitivity. Once these models encounter novel viewpoints, the alignment breaks down and they revert to texture-based decisions, exposing the fragility of their apparent shape bias. By contrast, viewpoint-diverse models show the reverse pattern: lower shape bias on familiar views but stronger reliance on shape under viewpoint distribution shift. This suggests that broader viewpoint diversity encourages networks to encode shape as a transferable property, enabling shape-based recognition to emerge specifically when generalization is required.

The Grad-CAM analyses revealed that viewpoint diversity stabilizes the spatial focus of recognition. Restricted-view models often dispersed their attention to background features under novel viewpoints, suggesting they lacked robust object representation that generalize across views, and might be leveraging background context or viewpoint-specific features (e.g. a particular edge or texture patch that was consistently seen in training images of that object; zhu2016object, seijdel2020depth). In contrast, viewpoint-diverse models consistently maintained object-centered attention, effectively segregating object from background even when confronted with novel viewpoints. This provides mechanistic evidence that viewpoint-diverse training induces a form of implicit scene segmentation, anchoring recognition to the object itself rather than background context Loke et al. (2024).

### Theoretical Implications

The contrast in learning dynamics suggests a natural interpretation through the lens of the bias–variance trade-off Geman et al. (1992). Viewpoint-restricted training produces low-variance, view-specific solutions that excel on familiar inputs but fail under distribution shift. By contrast, viewpoint-diverse training introduces greater variance during learning, which compels models to discover higher-level, more invariant features. In this sense, viewpoint diversity reshapes the trade-off: models sacrifice some in-distribution efficiency in exchange for representations that generalize more robustly. This pattern is mirrored in representational content. Shape-bias analyses reveal that reliance on shape is not a fixed network property but an adaptive strategy that emerges under conditions demanding generalization. Restricted-view models appear shape-biased on familiar views only because texture and shape cues are aligned, and viewpoint-diverse models choose shape cues when invariance is required.

Taken together, these findings provide an important clue about how inductive biases are shaped Lake et al. (2017). Rather than being hardwired, biases toward texture or shape emerge from the structure of visual experience. Viewpoint-restricted training fosters shortcut strategies tied to local regularities, while diverse training encourages networks to internalize shape as the more stable, transferable property.

### Connections to Human Vision

From a cognitive modeling perspective, these results provide a perspective that situates viewpoint diversity as a developmental driver of inductive bias, paralleling how human vision acquires a robust shape bias through exposure to varied perspectives early in life. Humans and other animals acquire viewpoint-invariant recognition through rich, early multi-view experience. Viewpoint-diverse models maintain object-centric attention, mirroring human visual processing that robustly tracks objects across viewpoint changes Lindsay (2020). Our work is inspired by findings that infants with opportunities to handle and visually inspect objects from many angles develop more viewpoint-invariant object perception than those with limited views Johnson (2010), and leveraged 3D object renderings to simulate this developmental principle for CNNs – essentially feeding the networks a more human-like “data diet”. This parallels recent work showing that toddler-like close-up views also reduce texture bias and increase human alignment Müller et al. (2025)

The outcome supports that when CNNs are trained under conditions more akin to human visual experience (rich, varied viewpoints and even active viewing), they develop more human-like robustness. Our results support that training with diverse viewpoints might counteract CNNs’ texture bias by compelling them to attend to more invariant shape features, which mirrors human development, where extensive exposure to varied perspectives fosters a stable preference for shape as the more invariant cue Landau et al. (1988); Diesendruck and Bloom (2003).

### Practical Implications

Toward human-like object perception, our study highlights that training data design is as crucial as network design for achieving human-like visual generalization. Modern CNN architectures certainly have the capacity for viewpoint invariance, as the same ResNet that fails on unusual viewpoints can learn to succeed when given the appropriate data Madan et al. (2022). The key is providing training experiences that emulate the variability of the real world. This means moving away from the “canonical viewpoint” bias of many curated datasets and toward viewpoint-diverse curricula. Our practical avenue is leveraging synthesized 3D data where one can systematically vary viewpoint, lighting, and context. Simulation allows generating unlimited images of an object from all angles, which is infeasible with manual photography. By integrating such synthetic multi-view data into model training, we can imbue CNNs with a more comprehensive understanding of objects. Standard data augmentation alone (e.g. adding noise or slight rotations to a restricted-view dataset) could not replicate this effect; while augmentation helps, it typically exposes the model to low-level perturbations around the same viewpoints, failing to develop true 3D invariances Alcorn et al. (2019). Our results add that explicitly incorporating viewpoint variation during training is another powerful way to achieve high-level feature learning that foster robustness to viewpoint change.

## Limitations

Our findings should be interpreted with several limitations in mind. First, our manipulation of viewpoint diversity relied on synthetic 3D renderings. Although this approach provides precise control over viewpoint and ground-truth segmentation, the models were not trained on natural photographs. Because synthetic imagery lacks the full complexity of real-world textures, lighting, and backgrounds, a domain gap may remain. While we expect the underlying mechanism we identify – namely, that broader viewpoint diversity encourages more invariant, shape-centered representations – to hold under naturalistic conditions, future work could address this domain gap by integrating more realistic datasets or employing complementary methods allowing photorealistic rendering of objects such as Neural Radiance Fields (NeRF, mildenhall2020nerf), Deep Signed Distance Functions (Deep SDF, park2019deepsdf) or Gaussian splatting Kerbl et al. (2023).

Second, while we studied static generalization to new views, another hallmark of human vision is the ability to actively seek information – e.g. a person who can’t recognize an object immediately might tilt their head or walk around the object until a familiar view comes into sight. It would be interesting to train embodied models or agents that can actively rotate objects (in simulation) during training, mirroring how toddlers handle toys Johnson (2010). Such active training might further improve viewpoint invariance and even efficiency of learning, as the agent could focus on the most informative viewpoints. Third, while our findings focus on convolutional neural networks, future work should examine whether similar benefits of viewpoint diversity extend to alternative architectures. For example, ViTs Dosovitskiy et al. (2021) and capsule networks Sabour et al. (2017) offer different inductive biases that may naturally support viewpoint generalization.

Finally, we focus on *category recognition*. However, viewpoint diversity may also benefit related tasks such as object detection, segmentation, or 3D pose estimation. Models trained with multi-view data may learn more complete object extents, improving localization and interpretation under perspective change. Beyond this, our analysis touched on texture-shape bias; future work could examine if viewpoint-diverse training can be combined with texture-debiasing methods (e.g., style transfer, geirhos2018imagenet) to further push CNNs toward human-like cue usage. Understanding the remaining differences between our best models and human vision is also important.

Viewed through a developmental lens, these limitations and extensions highlight the same principle: restricted-view training corresponds to an impoverished “data diet”, while child-like experience provides rich, varied, and actively sampled perspectives. Incorporating these conditions into artificial training regimes will move us closer to a truly human-aligned vision.

## Conclusion

Our study demonstrates that deliberately mimicking the viewpoint diversity of early human visual experience substantially improves viewpoint generalization in CNNs and brings their recognition strategies closer to those of humans. Models trained with diverse perspectives not only performed better under novel viewpoints, but also developed representations that favor shape over texture and maintain object-centered attention even on novel views. Next to architecture and training regime, we identify viewpoint diversity as a dimension that affects model-human alignment. Viewed through a developmental lens, this suggests that child-like visual experiences with rich in varied and actively sampled perspectives, point a direction for building more robust and human-aligned computer vision models.

## Usage of AI

The authors utilized ChatGPT to detect grammatical errors and reduce the number of words in the text. The authors have reviewed the article thoroughly and take full responsibility for the final version of this article.

## Supplementary Information

### Object Exemplars

**Figure S1.**
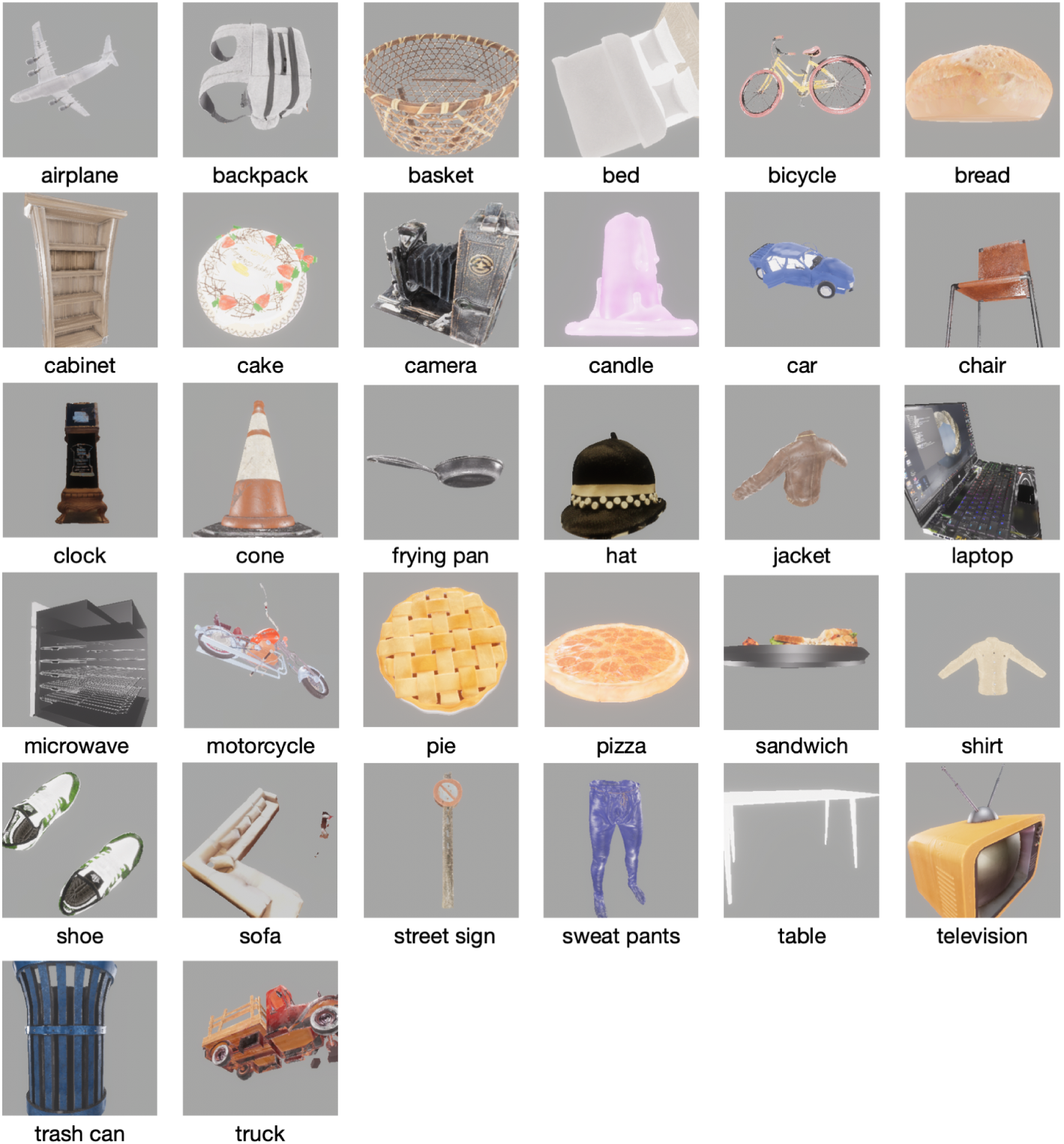
Dataset category exemplars. Those exemplars show the same 32 categories encompassed by every dataset generated in this study.

